# Cardiac belief updating from volatile physiological afferents

**DOI:** 10.64898/2026.09.04.749527

**Authors:** Nicolas Legrand, Lilian Weber, Christoph Mathys

## Abstract

Cardiac interoception is commonly assessed by comparing subjective estimates of heart rate with physiological measurements. But beliefs can easily obscure these measures, such that they cannot readily distinguish sensitivity to afferent signals from the influence of prior expectations; this ultimately challenges the notion that behaviours under these tasks could reflect interoceptive processes at all. Here, we develop a computational framework that uses naturally occurring heart rate variability to quantify how strongly perceptual beliefs are updated by physiological evidence. We formalise interoception as weighted Bayesian updating under the joint influence of expectations and afferents, and derive a new measure, cardiac interoceptive sensitivity, that quantifies the extent to which beliefs move with incoming signals. Applying this to the largest Heart Rate Discrimination dataset to date (n=549), a task providing robust estimates of cardiac beliefs, we find that sensitivity is weak in the healthy population, but shows large interindividual differences, with 43% of the participants exhibiting behaviours at least minimally compatible with interoceptive processing. These findings challenge the interpretation of conventional measures of cardiac interoception. They also introduce a new framework to relate bodily signals that cannot be controlled experimentally to external signals that can, laying a computational foundation for embodied psychophysics.

## Introduction

Cardiac interoception, the ability to sense signals arising from the heart, has been hypothesised to play a central role in regulating behaviour and shaping subjective experience **(Critchley & Garfinkel, 2017; Feldman et al., 2024; James, 1884; Khalsa et al., 2018)**. As psychology and computational neuroscience increasingly focus on individual differences in this domain, concerns have emerged regarding the reliability of experimental tasks and their associated computational metrics **(Desmedt et al., 2023)**. Central to these debates is the construct of interoceptive accuracy, the objective ability to detect and report cardiac signals **(Garfinkel et al., 2015)**. Yet task performance can arise from fundamentally different strategies: either accurately tracking ongoing physiological signals or relying on prior expectations only **(Desmedt & Bergh, 2024)**. Consequently, agreement between subjective reports and measured physiology does not, by itself, demonstrate that cardiac afferents shaped interoceptive behaviours **(Desmedt et al., 2018)**.

Measures of interoceptive accuracy have been facilitated by the ease of recording cardiac signals concurrently with behavioural reports, building on the idea that interoceptive judgments are informed by physiological signals **(Adamic et al., 2025; Epstein & Stein, 1974; Khalsa et al., 2009; Larsson et al., 2021; Mandler & Kahn, 1960; Pennebaker et al., 1982; Ponzo et al., 2021)**. This has underpinned now widely used paradigms, such as the heartbeat detection and discrimination tasks **(Flynn & Clemens, 1988; Savage et al., 2025)**, or the heartbeat counting (HBC) tasks **(Dale & Anderson, 1978; Schandry, 1981)**. But performance on these tasks can reflect heterogeneous strategies, including reliance on prior beliefs and generic knowledge. This limitation is most apparent in the HBC task, where the influence of prior expectations rather than physiological tracking has been repeatedly documented **(Desmedt et al., 2018; Desmedt et al., 2020; Ring & Brener, 1996; Ring & Brener, 2018)**.

The Heart Rate Discrimination (HRD) task **(Legrand et al., 2022)** focuses instead on the bias and precision of beliefs themself, accounting for their predominance in interoceptive judgments. The task requires participants to compare their perceived heart rate with an external auditory reference, where the bias is manipulated by a staircase procedure **(Kontsevich & Tyler, 1999)**. Psychometric analyses yield estimates of threshold (biases) and slope (square root of the inverse precision) as well as metacognitive sensitivity derived from confidence ratings **(Fleming, 2017)**. The task has been extensively studied **(Banellis et al., 2026a; Banellis et al., 2026b)**, consolidated in its methods **(Courtin et al., 2025)** and adapted in a variety of contexts, including meditation **(Palmer et al., 2025; Rudnicki et al., 2025)**, drug studies **(Leganes-Fonteneau et al., 2025; Tyrer et al., 2025)**, or clinical populations **(Grondel et al., 2026; Jeganathan et al., 2024)**.

But to date, the computational models available **(Courtin et al., 2025; Legrand et al., 2022)** covertly assume that the referent signal (heart rate) is invariant when estimating beliefs and their precision (see Section 11.1 for a formal demonstration). These tools are excellent for characterizing what participants think about their heart rate over a fixed period of time, but not how beliefs are coupled with physiological afferents, which is closer to interoceptive processing as already explored in the respiratory domain **(Brand et al., 2024; Harrison et al., 2021)**. In particular, they do not distinguish whether beliefs are dynamically updated or remain largely insensitive to physiological afferents. This fork in multiple strategies, in turn, introduces confounds when paired with a volatile reference signal, which is what the human heart produces, as cardiac variability can directly influence precision estimates.

Here, we derive a new computational metric: interoceptive sensitivity, which is the extent to which cardiac beliefs change with incoming evidence. We suggest that it lies along a continuum between two limiting strategies: 1) either disregarding afferent signals, or relying on them exclusively (see Figure 1). While measuring this influence would normally require the rigorous manipulation of the two signals of interest, we demonstrate that naturally occurring heart rate variability can partially replace experimental control and provide a window on the causal influence of physiological signals.

**Figure 1:**
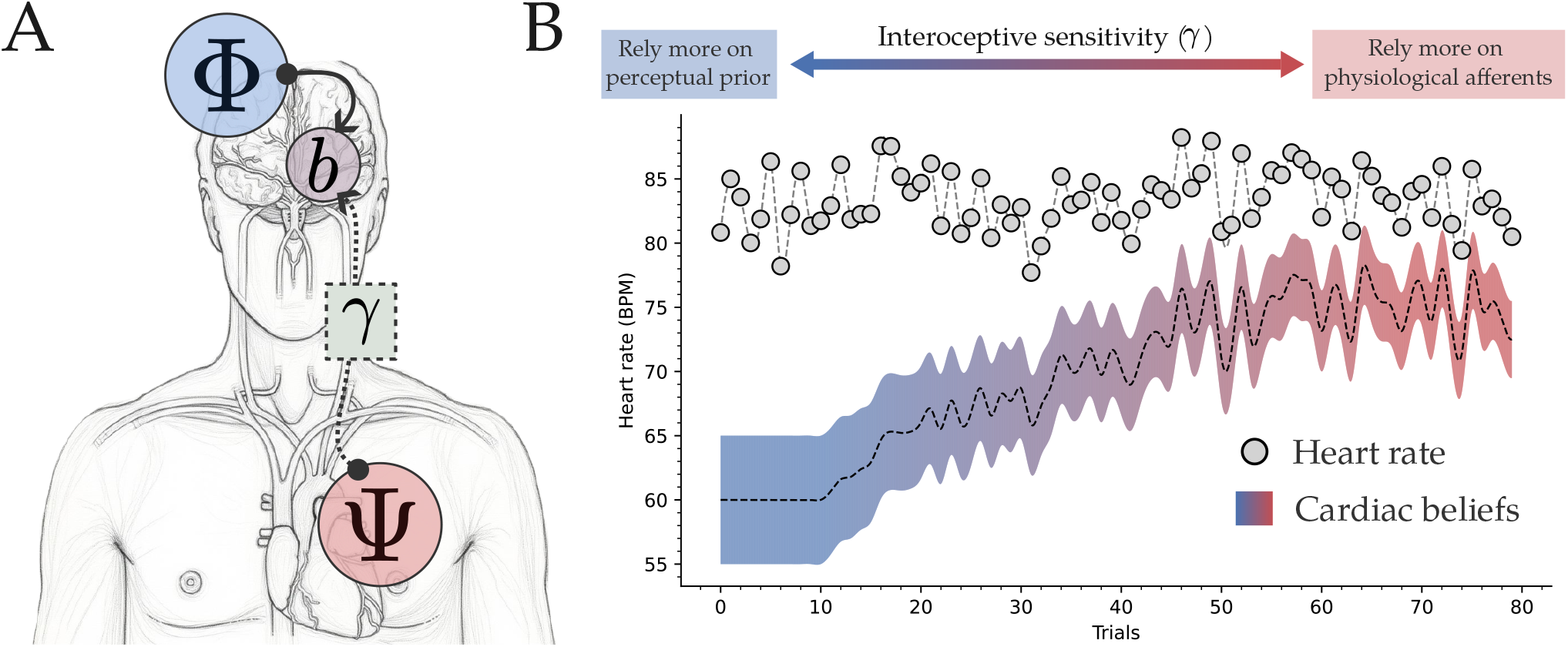
Cardiac belief updating along the sensitivity axis **A**. Schematic representation of the two sources of information influencing cardiac interoceptive beliefs (*b*), from prior perceptual expectation (Ψ) and physiological afferents (Φ). Interoceptive sensitivity weights the influence of afferent signals on the belief updates. **B**. Simulated belief trajectories with increasing levels of interoceptive sensitivity. The grey dots and lines represent instantaneous heart rates, which can be derived from RR intervals or averaged over arbitrary time windows. The cardiac beliefs and their precision are represented with a gradient of shaded areas, from no sensitivity (*γ* = 0.0, blue) to the highest sensitivity (*γ* = 1.0, red). Higher sensitivity (*γ*) increases the influence of physiological afferents and results in covarying beliefs and physiology. When *γ* = 0, cardiac beliefs are only guided by perceptual priors. When *γ* = 1.0, cardiac beliefs strictly covary with physiological afferents. Note that bias and uncertainty can remain, in theory, even under perfect sensitivity.

The paper provides the computational backbone to infer cardiac interoceptive sensitivity from the Heart Rate Discrimination task **(Legrand et al., 2022)**. We formalise sensitivity as the degree to which Bayesian beliefs incorporate cardiac afferents over time. We introduce two generative models with static and dynamic priors to estimate the weight assigned to physiological input during belief updating. Using the largest HRD dataset to date (n = 549), we validate our approach and show that cardiac afferents weakly influence healthy participants, yet exhibit substantial interindividual variability. This framework provides a principled measure of interoceptive sensitivity and a generic modelling approach for embodied psychophysics, where researchers seek to relate bodily signals that cannot be controlled experimentally to external signals that can.

## Materials and methods

### Experimental design

We re-analyse the largest HRD task dataset (n=549) previously published in **Legrand et al. (2022)** and **Banellis et al. (2026a)**. For details regarding the experimental design, the reader should refer to **Legrand et al. (2022)**. The dataset consists of two sessions (n=218 for session 1 and n=518 for session 2, with n=187 who completed both sessions). Here, we report results on the second session, and we use the two sessions together for the retest analysis. In a nutshell, participants are asked to discriminate the frequency of their heart rate and an external tone frequency for reference. We distinguish **1)** a perceptive phase, where the participant focuses on a reference, either the heart rate (i.e., interoception) or an external tone (i.e., exteroception), followed by **2)** a discrimination phase, where another external tone is presented, and the participant should indicate if it is faster or slower than the reference. A Bayesian staircase **(Kontsevich & Tyler, 1999)** is then applied to manipulate the bias value (*α*) that represents the difference in frequency between the reference, either the heart rate (interoception) or the first tone (exteroception), and the second tone. By producing decisions repeatedly (i.e., is the second tone faster or slower?), it is possible to infer the threshold (i.e., bias) and slope (i.e., square root of the inverse precision) of the underlying psychometric function.

### Computational models

We evaluated four computational models of cardiac interoception that differed in how cardiac beliefs incorporate physiological information across trials. We first compared two limiting hypotheses. Under the **dynamic** model, cardiac beliefs track physiological fluctuations from trial to trial. Under the **static model**, cardiac beliefs remain stable despite fluctuations in heart rate. The remaining generative models interpolate between these two extremes. Specifically, we considered two Bayesian models. The first performs a **weighted Bayesian update** from a fixed prior independently at each trial. The second extends this formulation by allowing posterior beliefs to become priors for subsequent trials, thereby introducing temporal dependencies. This last model is implemented as a generalised hierarchical Gaussian filter (HGF) **(Mathys et al., 2011; Mathys et al., 2014; Weber et al., 2026)**, which we refer to as the **cardiac HGF**.

### Mathematical notations

Normal distributions are parametrised by their mean (*μ*) and variance (*σ*^2^). The precision, sometimes used for convenience, is denoted *π* and is given by 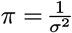. The cumulative distribution function of a normal distribution is given by:

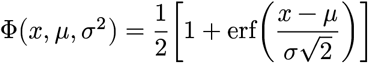

With erf(⋅) denoting the error function.

The task consisted of *N*_*e*_ exteroceptive trials and *N*_*i*_ interoceptive trials. *N* was set to 80 in the first session in **Legrand et al. (2022)**, and set to 60 otherwise. Each exteroceptive trial required the participant to discriminate the highest frequency between two tones. We denote 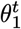 and 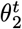 as the frequencies of the first (reference) and second (decision) tone, respectively.

Similarly, each interoceptive trial required the participant to discriminate the higher frequency between the objective heart rate *h*^*t*^ (reference), measured by the pulse oximeter, and another tone 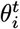 (decision). For modelling convenience, trial-wise heart rates are treated as samples from a participant-specific distribution:

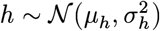

This variable *h*, which is directly observed, should be distinguished from the cardiac belief *b*, which is an unobserved latent variable inferred from behaviour. For convenience, we consider that this cardiac belief is also normally distributed:

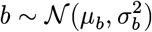

A similar distinction can be made between the objective tone frequency and the subjective tone frequency, which is omitted here for concision.

Participants should then assess whether the second tone was faster than what they infer from the reference signal 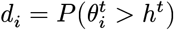 (i.e., interoception) or the first tone 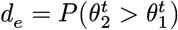 (i.e., exteroception). Here (*d*) represents the Bernoulli probability that the tone is faster than the reference.

### Model 1: dynamic beliefs

The **dynamic belief** model assumes that cardiac beliefs track physiological fluctuations on every trial. Consequently, the latent cardiac belief covaries with the observed heart rate up to a participant-specific bias and sensory uncertainty. This corresponds to the psychometric model introduced in the original HRD task **(Legrand et al., 2022)**, with subsequent extensions differing only in the inclusion of lapse parameters **(Banellis et al., 2026a; Banellis et al., 2026b; Courtin et al., 2025; Tyrer et al., 2025)**. The model assumes cardiac belief *b*^*t*^ covaries with the objective heart rate *h*^*t*^, with a bias *α* and some noise *β*. At each trial, the probability of responding “Faster” is given by:

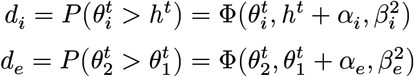

For interoceptive and exteroceptive trials, respectively. The decision variance reflects uncertainty in both the comparison tone and the internal reference. Assuming constant auditory variance 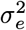 across trials gives

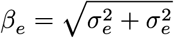

for exteroception and

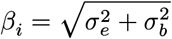

for interoception.

### Model 2: static beliefs

The **static beliefs** model assumes that cardiac beliefs *b*^*t*^ are not influenced by the objective heart rate *h*^*t*^. The probability of responding “Faster” at each trial is then given by:

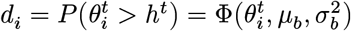

This is equivalent to fitting the psychometric function on the frequency of the second tones only, indicating that the participants compare the auditory stimulus with a stable internal representation of heart rate. The **dynamic** and the **static** models therefore define two limiting hypotheses regarding the contribution of cardiac afferents to perceptual decisions.

### Model 3: Weighted Bayesian update with a static prior

It is also possible to assume continuous policies that would range between these two extremes. In the **weighted updates**, cardiac beliefs are modelled as the Bayesian combination of a fixed prior and the current afferent observation. The prior is represented by 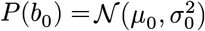, whereas the afferent evidence at trial *t* follows 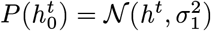. A weighting parameter 0 ≤ *γ* ≤ 1 determines the relative influence of afferent evidence during the update. The resulting posterior precision and mean are

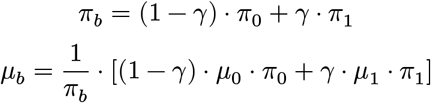

Although *γ* controls the nominal weighting of afferent evidence, its effective contribution also depends on the relative precisions of the prior 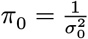 and sensory observation 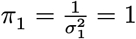. The precision of the afferent observation is fixed to one, defining the scale of the latent precision parameters and preventing non-identifiability. Consequently, *γ* alone is not directly interpretable as interoceptive sensitivity. We define interoceptive sensitivity as the relative contribution of afferent evidence to the posterior precision

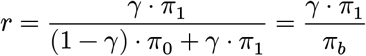

and express this quantity on the logit scale,

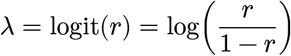

where *λ* = 0 indicates equal contributions of prior expectations and afferent evidence, positive values indicate greater reliance on afferent evidence, and negative values indicate greater reliance on prior expectations.

### Model 4: Weighted Bayesian update with a dynamic prior

The previous model assumes a fixed perceptual prior throughout the task. We next relax this assumption by allowing posterior beliefs to become priors on subsequent trials. This introduces temporal dependencies in cardiac beliefs and enables gradual adaptation to sustained physiological changes. We implement this process using a generalised Hierarchical Gaussian Filter (HGF) **(Weber et al., 2026)**, a Bayesian state-space model in which belief updates are governed by precision-weighted prediction errors **(Mathys et al., 2011; Mathys et al., 2014)**.

Separate but structurally identical HGFs were fitted to the interoceptive and exteroceptive conditions (see Figure 4 A.). In each network, lower-level nodes encode noisy sensory observations, whereas higher-level nodes represent latent beliefs about the corresponding stimulus. Beliefs are updated sequentially across trials through precision-weighted prediction errors. Following HGF terminology, higher-level latent states are referred to as value parents **(Weber et al., 2026)** because they specify the expected value of lower-level observations.

In the exteroceptive network, two continuous input nodes *x*_1_ and *y*_1_ receive the first and second tone frequencies (respectively), parametrised as follows:

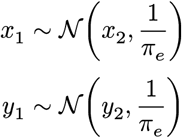

Where *π*_*e*_ is the precision of the auditory prior, assuming that this quantity is constant between reference and decision phases. Both nodes are influenced by their value parents *x*_2_ and *y*_2_.

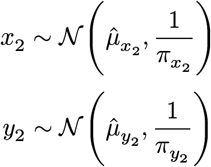

An auditory prior *μ*_*e*_ is used as the initial state for both 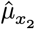 and 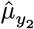; afterwards, this value is inherited from previous updates.

Learning is governed by two complementary mechanisms. Tonic volatility controls how rapidly latent beliefs are allowed to evolve across trials, whereas sensory precision determines the confidence assigned to incoming observations. Because these parameters influence different components of the update, participants may have imprecise and rapidly shifting cardiac beliefs while remaining perceptually certain, or conversely maintain highly precise cardiac beliefs while being perceptually uncertain (see examples of this behavior in Figure 4, C.).

The learning rate is controlled by *ω*_*e*_, the tonic volatility of the auditory beliefs, a fixed parameter that does not evolve across time. Auditory judgments should be characterised by large tonic volatilities (i.e., current auditory belief is mainly influenced by the incoming signal), as participants are usually able to discriminate tones with only 1 or 2 bpm deviations **(Legrand et al., 2022)**.

Importantly, here, the precision of the auditory prior *π*_*e*_ is part of the decision function; therefore, while both parameters compete with each other, we are not facing a case of non-identifiability. The decisions to answer “Faster” or “Slower” are generated from the posterior distributions inferred by the HGF and reflect the subjective confidence that *y*_2_ is greater than *x*_2_. This is then equivalent to computing the distribution of the differences *D*:

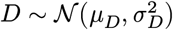

with:

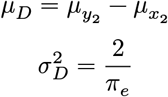

and to compare it to 0:

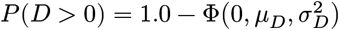

A similar principle is applied to interoceptive decision, with the difference that the auditory precision is shared with the exteroceptive network, allowing us to model cardiac and auditory precision separately. Cardiac signals are received through the continuous input node *x*_1_. This node is itself the value child of the continuous node *x*_2_ that is encoding the cardiac belief, such that:

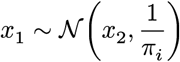

and, as before, for the reference tone:

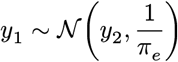

Because HGF updates are themselves Bayesian precision-weighted updates, the effective contribution of sensory evidence can be expressed on the same scale as in Model 3.

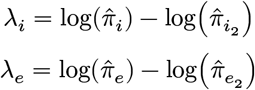

Where 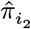 and 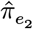 are the expected precision of the value parent for the interoceptive and exteroceptive branches, respectively. We provide the full derivation of this equality in Section 11.2.

### Software and statistical analyses

The computational models reported here are based on the generalised hierarchical Gaussian filter **(Weber et al., 2026)** as implemented in PyHGF v0.3.0 **(Legrand et al., 2026)**. We used Hamiltonian MonteCarlo (NUTS sampler) **(Betancourt, 2017; Hoffman & Gelman, 2011)** as implemented in PyMC v6.0.1 **(Abril-Pla et al., 2023)** and Nutpie v0.16.10 to approximate Bayesian inference and recover parameters from observed behaviors. Model comparison was performed using the expected log pointwise predictive density (ELPD) **(Vehtari et al., 2016)** on the interoceptive trials, as implemented in Arviz v1.2.0 **(Kumar et al., 2019)**. We estimated the ELPD using Pareto-smoothed importance sampling leave-one-out cross-validation (LOO).

Statistical testing was performed using Pingouin v0.6.1 **(Vallat, 2018)**. Due to the presence of outliers in some of the variables of interest, we assessed the robustness of correlations for test-retest reliability using skipped correlations **(Pernet et al., 2013)**. Discriminability between task modalities was quantified as the common-language effect size (CLES), that is, the probability that a randomly drawn exteroceptive estimate exceeds a randomly drawn interoceptive one. Effect sizes were obtained with Pingouin **(Vallat, 2018)**, with 95% confidence intervals from 5,000 bootstrap resamples and significance assessed by Mann–Whitney U tests treating the two modalities as independent samples. We used Matplotlib **(Hunter, 2007)** and Seaborn **(Waskom, 2021)** for data visualization.

The code used to produce the simulations and analyse experimental data is available at https://github.com/LegrandNico/ComputationalCardioception. The dataset analysed in this paper was obtained using the Cardioception toolbox **(Legrand et al., 2022)**.

## Results

### Static cardiac beliefs better explain HRD performances

We first compared two competing hypotheses of cardiac belief formation. The dynamic-belief model suggests that participants continuously update their representation from trial-to-trial heart rate fluctuations, while the static-belief model suggests that this representation remains stable despite physiological variability (Figure 2 panel A). We compared the psychometric fits obtained from each model (Figure 2D) using Bayesian model comparison. The leave-one-out expected log pointwise predictive density (ELPD) showed that the static-belief model provided a better account of behaviour for most participants (Figure 2B). Of the 518 participants included in this analysis, 421 (81%) had their performances better explained by a model of static beliefs; the difference was larger than the uncertainty (the SE estimate did not include 0) for 288 of them (55%). By contrast, only 97 participants (18%) had their performances better explained by a model of dynamic beliefs, and this was larger than uncertainty for 25 of them (5%). These results indicate that, when directly contrasted, models assuming stable cardiac beliefs explain HRD performance more often than models assuming trial-by-trial tracking of cardiac physiology.

**Figure 2:**
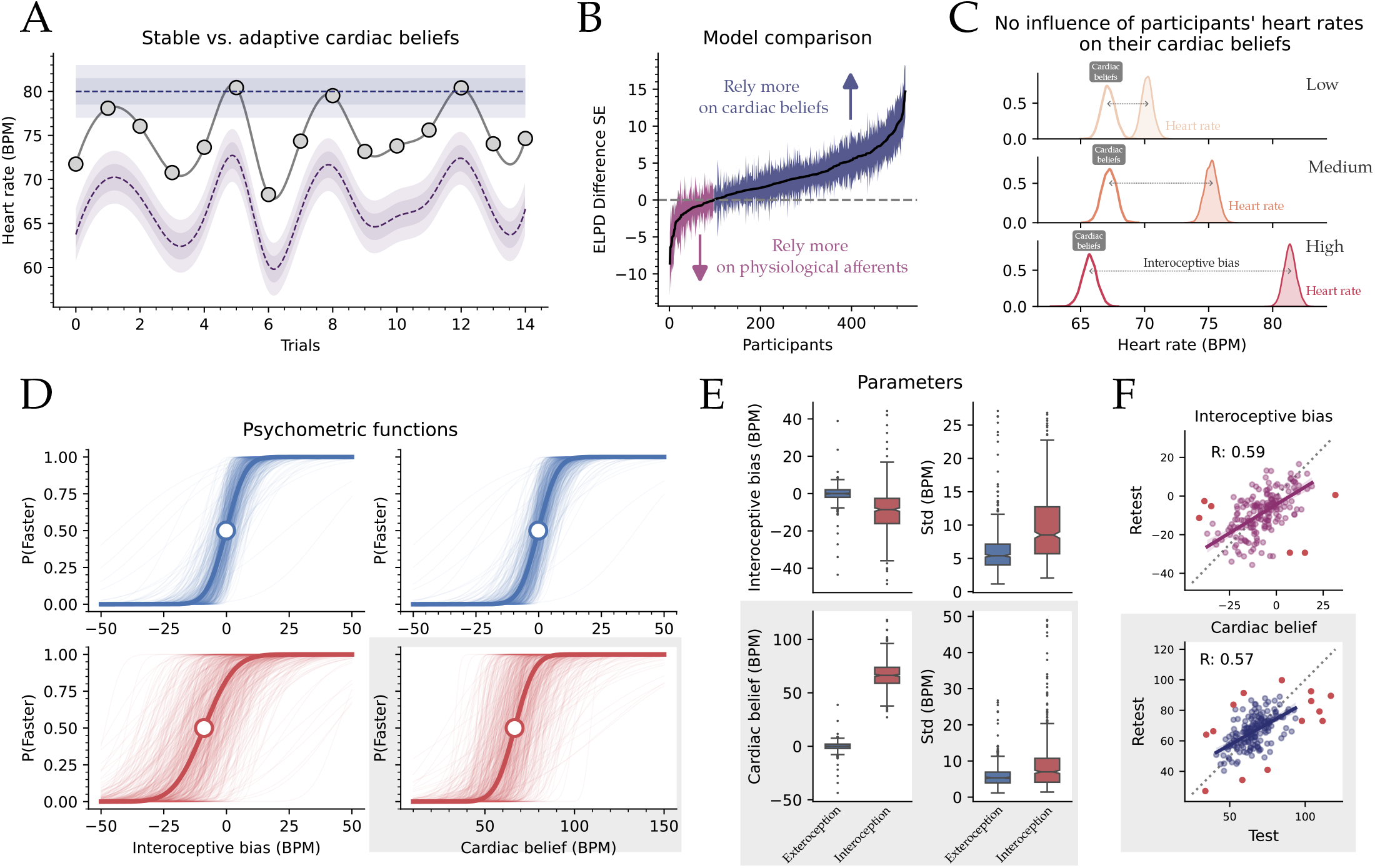
Behaviors at the HRD task are better explained by static cardiac beliefs. **A**. Simulation of instantaneous heart rate (grey dots and line), static beliefs (blue line and shaded areas), and dynamic beliefs (purple line and shaded area). These two models represent the two extremes of interoceptive sensitivity, where prior expectations can have an all-or-none influence. **B**. Model comparison between static (blue) and dynamic (purple) cardiac beliefs. Static beliefs better explain the behaviors among a majority of participants. **C**. Cardiac beliefs are stable, despite significant fluctuation in the underlying heart rates. By splitting the trials using the two tertiles of the subject’s heart rate, we observed that cardiac beliefs (transparent distributions) are stable compared to the objective heart rates (shaded distributions). As heart rate fluctuates across low (yellow), medium (orange), and high (red) frequencies, beliefs hold and bias increases. **D**. Psychometric fit along the bias for dynamic belief (left panel) and heart rate for static beliefs (right panel). The exteroceptive trials are fitted using a dynamic model in both cases in order to learn the bias; the functions are therefore near-identical. **E**. Distribution of psychometric parameter for static (lower panel) and dynamic beliefs (upper panel). Similarly, cardiac beliefs and exteroceptive biases in the lower left panel are coming from two different models and should not be compared statistically. **F**. Test-retest reliability for the dynamic (purple, upper panel) and static models of beliefs (blue, lower panel).

That observation was made by assuming a fixed strategy and a fixed heart rate generation across the trials. We therefore next asked whether cardiac beliefs would remain stable across naturally occurring changes in heart rate. Because heart rate could not be experimentally manipulated as in other studies **Windmann et al. (1999)**, we grouped each participant’s trials into low-, medium-, and high-heart-rate tertiles and estimated cardiac beliefs separately for each subset (Figure 2C). Here, despite clear differences in physiological heart rate across tertiles (low: *μ* = 70.21, *σ* = 0.45, ETI_89_ = [69.39, 71.09], medium: *μ* = 75.20, *σ* = 0.46, ETI_89_ = [74.31, 76.06], and high: *μ* = 81.36, *σ* = 0.46, ETI_89_ = [80.52, 82.22]), inferred cardiac beliefs remained remarkably stables (low: (*μ* = 67.14, *σ* = 0.54, ETI_89_ = [66.13, 68.14]), medium: *μ* = 67.24, *σ* = 0.55, ETI_89_ = [66.28, 68.35], and high: *μ* = 65.71, *σ* = 0.56, ETI_89_ = [64.60, 66.69]). Thus, substantial fluctuations in physiological state were accompanied by little to no change in participants’ inferred cardiac beliefs.

The recovered psychometric parameters (Figure 2D, E) reproduced the characteristic underestimation of heart rate previously reported for the HRD task **(Legrand et al., 2022)**. Interoceptive thresholds were lower than exteroceptive thresholds (*μ*_Intero_ = −8.57, *μ*_Extero_ = −0.16, CI_95%_ = [7.76, 9.83], *t*_517_ = 16.71, *p* < 0.001, BF_10_ = 9.51*e* + 46, *d* = 0.96). Interoceptive slopes were also larger than exteroceptive slopes both under the dynamic-belief model (*μ*_Intero_ = 9.55, *μ*_Extero_ = 6.07, CI_95%_ = [−3.99, −2.97], *t*_517_ = −13.45, *p* > 0.001, BF_10_ = 1.65*e* + 32, *d* = 0.79), and under the static-belief model (*μ*_Intero_ = 8.77, *μ*_Extero_ = 5.93, CI_95%_ = [−3.44, −2.22], *t*_517_ = −9.13, *p* > 0.001, BF_10_ = 2.61*e* + 15, *d* = 0.50).

### Weighted Bayesian updating reveals graded interoceptive sensitivity

Because neither limiting model accounted for all participants, we next asked whether cardiac belief updating is better described as a continuum between a stable prior and fully dynamic beliefs. We therefore estimated interoceptive sensitivity (*λ*), which quantifies the contribution of afferent cardiac signals to Bayesian belief updating (see Figure 3 A.).

**Figure 3:**
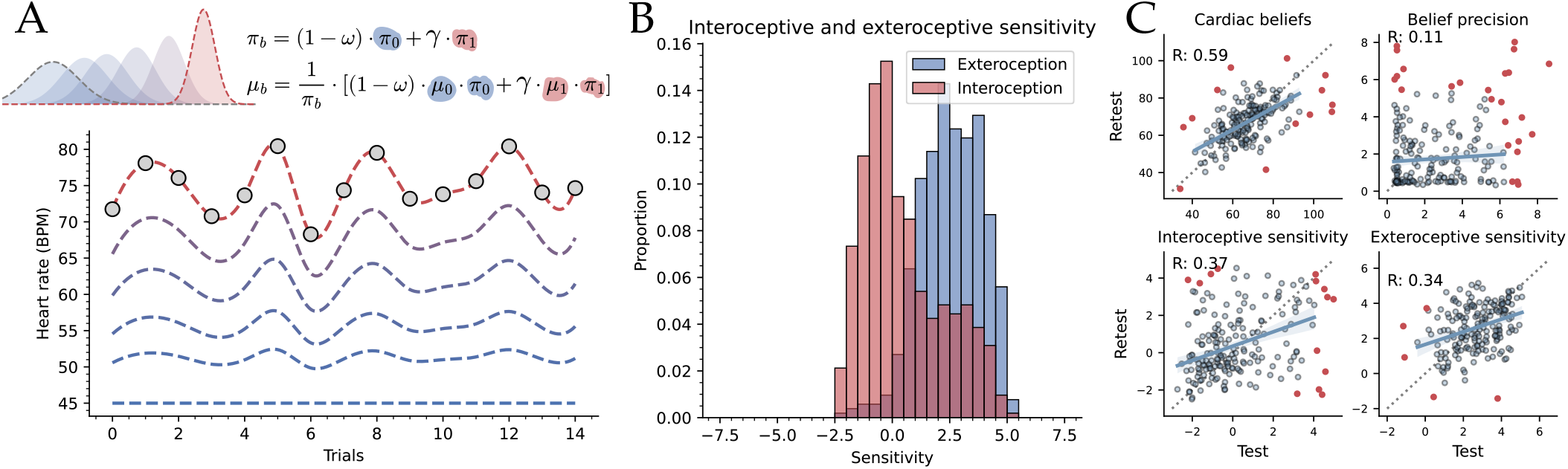
Cardiac beliefs as a weighted Bayesian update. **A**. Illustration of belief trajectory dynamics as a function of interoceptive sensitivity (*λ*), the logit-transformed weight of a Bayesian update (*γ*). Here, the update is applied without time dependence, such that the prior expectation is fixed and shared across trials. **B**. Discriminability of task modality using interoceptive sensitivity scores. **C**. Test-retest reliability for key metrics of interest.

Sensitivity differed sharply between modalities (see Figure 3 B.). Exteroceptive sensitivity was high (*λ*_*e*_: M = 2.56, SD = 1.33), corresponding to a weight of 0.88 (SD = 0.14) on the tone. By contrast, interoceptive sensitivity fell close to the point of equal contribution (*λ*_*i*_: M = 0.49, SD = 1.75; weight 0.55, SD = 0.29). The difference of 2.07 logits was large and highly reliable (*t*_517_ = 22.67, *p* < 0.001, CI_95%_[1.89, 2.25], *d* = 1.33, BF_10_ = 7.1*e* + 75), indicating that cardiac beliefs rely substantially less on incoming sensory evidence than auditory perceptual judgments.

Interindividual variability was also larger in the interoceptive domain: *λ*_*i*_ spanned −2.50 to 5.16, and its median (−0.02) fell below its mean, indicating that a majority of participants weighted the afferent signal at or below equal contribution while a minority weighted it heavily. Using modality as the class and model parameters as predictors, interoceptive sensitivity separated interoceptive from exteroceptive participant-level parameter estimates at AUC = 0.817, and the precision of the inferred belief at AUC = 0.939, both above the slope of the static-belief model (AUC = 0.624) and the threshold of the dynamic-belief model (AUC = 0.795).

Finally, across participants tested twice (*n* = 187, skipped correlations), the inferred cardiac belief was the most reliable parameter (*r* = 0.59), followed by interoceptive sensitivity (*r* = 0.37). The precision of the cardiac belief did not replicate (*r* = 0.11). Thus, Bayesian belief updating provides a measure of interoceptive sensitivity that captures meaningful individual differences while retaining moderate temporal stability.

### The cardiac hierarchical Gaussian filter captures dynamic belief updating

The weighted update treats the perceptual prior as fixed across the task. Relaxing this assumption lets the prior itself carry information forward, so that a belief can drift towards the afferent signal. We implemented this using a generalised hierarchical Gaussian filter **(Legrand et al., 2026; Weber et al., 2026)** with two value-coupled networks, one per modality (see Figure 4 A.).

Simulations confirmed that the model spans the continuum between static and dynamic belief updating (see Figure 4 C.). With low input precision and negative tonic volatility (i.e., low learning rate from the second level), the belief remains flat and wide, reproducing static beliefs; with high input precision and positive tonic volatility (i.e., high learning rate), it tracks the heart rate trial by trial, reproducing dynamic beliefs.

Fitted to behaviour (*n* = 518), the model reproduced and strengthened the modality difference observed in the weighted update. Exteroceptive sensitivity was uniformly high (*λ*_*e*_: M = 24.25, SD = 3.04, median = 24.72), whereas interoceptive sensitivity was negative in most participants (*λ*_*i*_: M = −1.85, SD = 6.17, median = −4.29), indicating that the afferent signal contributed less posterior precision than the prior (*t*_517_ = 81.42, *p* < 0.001, CI_95%_[25.47, 26.73], *d* = 5.37, BF_10_ = 4.7*e* + 292).

The *λ* parameter ranges to more extreme values in this model compared to the weighted update because a dynamic prior allows the ratio of afferent to prior precision to grow without the bound imposed by a fixed prior. As a result, exteroceptive sensitivity is close to its ceiling in almost every participant, suggesting that auditory judgements are driven almost entirely by the stimulus, and not as a graded individual measure. This enhanced separation translated into near-perfect discrimination between modalities. Using sensitivity alone, our model classified interoceptive and exteroceptive conditions with an ROC-AUC of AUC = 0.991 (see Figure 4 B. and Figure 5 C.). This suggests that the parameter is highly sensitive to the distinction between modalities and accentuates the computational distinction between modalities.

**Figure 4:**
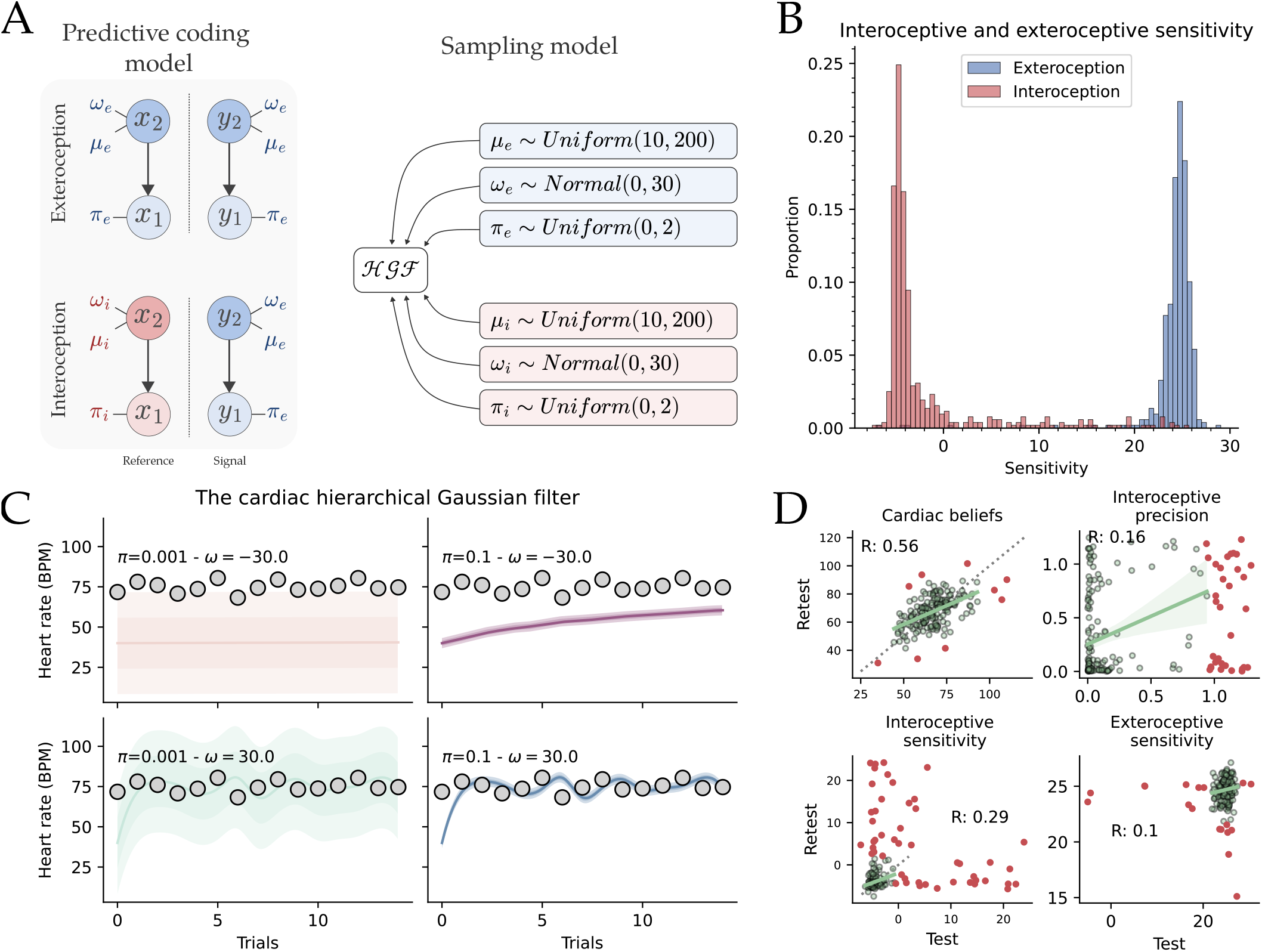
Overview of the cardiac hierarchical Gaussian filter (HGF). **A**. Schematic representation of the predictive coding model (left) and the graphical model used for sampling. Interoceptive and exteroceptive trials are handled by the lower and upper parts of the network (respectively). For each reference and decision tones/heart rate, a pair of observation nodes and a value parent guide the belief update. **B**. The cardiac HGF can discriminate task modality with close-to-perfect performance. **C**. Illustration of the influence of tonic volatility (*ω*) and input precision on the belief trajectories. Shaded areas represent ± 0.5 and 1.0 standard deviations around the mean. **D**. Test-retest reliability of some parameter of interest. This model can capture cardiac belief with performance comparable to both the static belief and weighted Bayesian updates. Red points represent outliers detected by the skipped correlation procedure.

**Figure 5:**
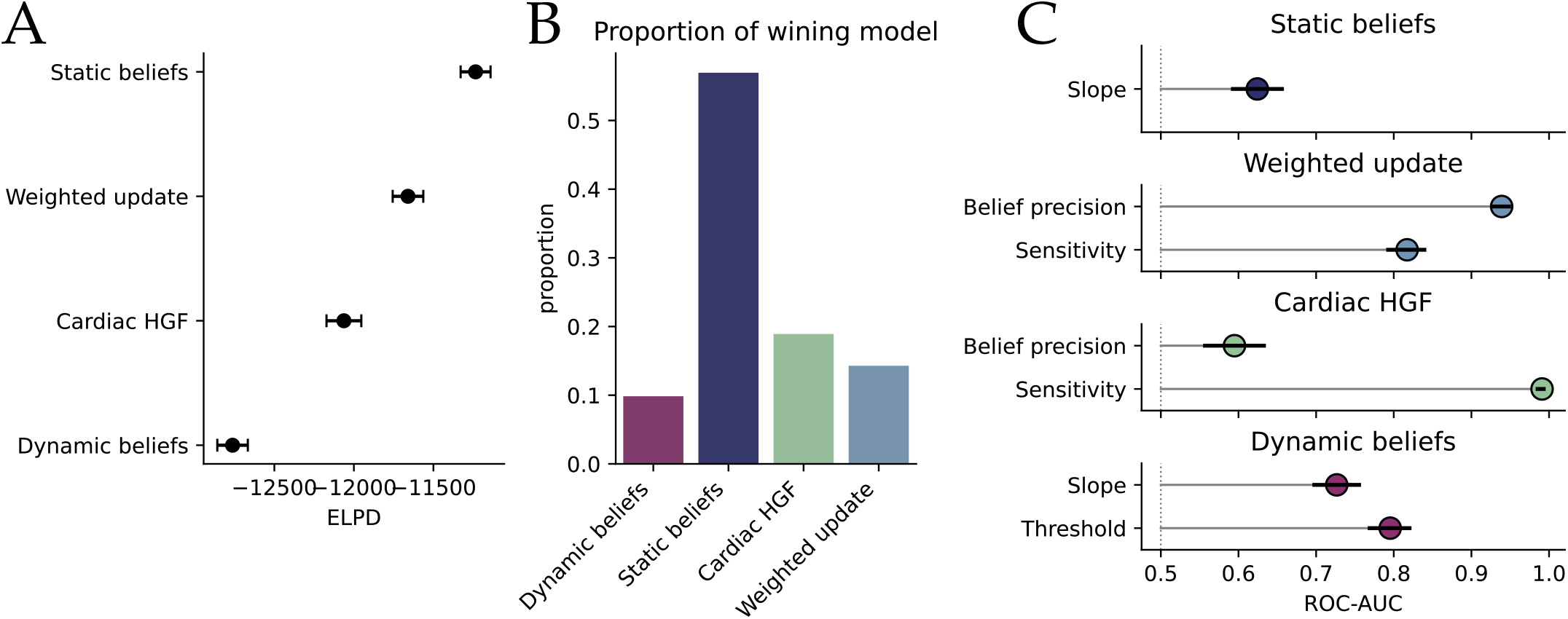
Comparison of four models of cardiac interoception. **A**. Model performance assessed by their expected log pointwise predictive density (ELPD) **(Kumar et al., 2019; Vehtari et al., 2016)**. A model assuming dynamic beliefs was the least efficient, while a model assuming static beliefs was the most efficient, in coherence with what is reported in Figure 2B. Models assuming Bayesian belief updates had intermediate scores, with the static priors eliciting better performance. **B**. Proportion of winning models at the individual level. Static beliefs were the best models for more than half of the participants, but the two Bayesian updates accounted for 14% and 19% of the participants each. Dynamic beliefs could explain only 9.8% of the population’s behavior. **C**. Discriminability of task modality from each model’s parameters, as the rank-based area under the ROC curve with 95% bootstrap confidence intervals over participants (5,000 resamples).

Across participants tested twice (*n* = 187), inferred cardiac belief remained moderately reliable (*r* = 0.56), whereas interoceptive precision showed weaker reliability (*r* = 0.16). By contrast, neither interoceptive sensitivity (*r* = 0.29, raw Pearson *r* = 0.09, CI_95%_ = [−0.06, 0.23]) nor exteroceptive sensitivity (*r* = 0.12, raw Pearson *r* = −0.02) showed strong test–retest reliability. Thus, although the HGF provides the clearest computational dissociation between modalities, this increased discriminability is accompanied by reduced stability of individual sensitivity estimates.

### Comparing models of cardiac interoception

Finally, we wanted to compare the four computational models across predictive performance, individual model preference, discriminability between task modalities, and test–retest reliability to evaluate their respective strengths.

At the group level, the static-belief model achieved the highest predictive performance, outperforming both Bayesian updating models and the original dynamic-belief model (see Figure 5A). This last model was used in the original HRD publication **(Legrand et al., 2022)** and extended in follow-up studies **(Banellis et al., 2026a; Banellis et al., 2026b; Courtin et al., 2025)**, but scored last on this comparison.

Model preference varied across individuals but showed a similar pattern (see Figure 5B). The static-belief model offered the best fit for a majority of participants (295, 57%), while weighted Bayesian updates and the cardiac HGF were preferred by 14% and 19%, respectively. Only 9.8% of participants were best described by the dynamic-belief model. Thus, although stable cardiac beliefs predominated, dynamic and Bayesian updating models captured substantial variation across individuals (43% of the population could be described as interoceptors according to this description).

The models differed more markedly in their ability to distinguish interoceptive from exteroceptive processing (see Figure 5C). The cardiac HGF achieved near-perfect discrimination using its sensitivity estimate alone (AUC = 0.991), outperforming the weighted Bayesian update using sensitivity and belief precision (AUC = 0.817 and AUC = 0.939, respectively). The threshold parameter of the dynamic-belief model also discriminated between modalities (AUC = 0.795), similar to the slope (AUC = 0.727), reflecting the documented underestimation of heart rate observed during interoception. By contrast, the precision parameter of the static-belief model provided little discriminative information (AUC = 0.624), and similar results were observed with the belief precisions in the HGF model (AUC = 0.595).

Finally, we compared the temporal stability of the inferred parameters across repeated testing (see **Legrand et al. (2022)** for details), but this time also including the static beliefs psychophysics. We found a medium reliability in interoceptive bias (*r* = 0.59, CI_95%_ = [0.49, 0.68], *p* < 0.001, see Figure 2 F, higher panel) and a medium reliability in cardiac belief (*r* = 0.57, CI_95%_ = [0.46, 0.67], *p* < 0.001, see Figure 2 F, lower panel). While Bayesian models recovered comparable levels of reliability in their estimates of cardiac beliefs, the measure of sensory precision and sensitivity did not elicit good test-retest reliability.

## Discussion

Cardiac interoception has long been challenged by the difficulty of disentangling the influence of physiological afferents from that of prior beliefs on perceptual decisions **(Desmedt et al., 2020; Ring & Brener, 1996; Ring & Brener, 2018)**. Here, we introduce a computational definition of cardiac interoceptive sensitivity: rather than asking whether participants can estimate their heart rate, our framework asks how strongly their beliefs are updated by changes in cardiac physiology. Using behavioural data from the largest Heart Rate Discrimination (HRD) dataset to date **(Banellis et al., 2026a; Legrand et al., 2022)**, we developed computational frameworks that exploit naturally occurring cardiac variability to estimate this quantity. We observed that beliefs in healthy participants were influenced only weakly by moment-to-moment physiological fluctuations compared with exteroceptive judgments, although substantial interindividual variability emerged. These findings distinguish stable cardiac beliefs from dynamic physiological tracking and provide a computational framework for quantifying this quantity in embodied psychophysics experiments.

### Cardiac interoceptive beliefs are predominantly static

Cardiac interoception is commonly evaluated in terms of accuracy, defined as the correspondence between subjective reports and physiological states **(Garfinkel et al., 2015)**. In tasks such as the HRD **(Legrand et al., 2022)** and heartbeat counting **(Dale & Anderson, 1978; Schandry, 1981)**, this correspondence is typically quantified as a bias score. But bias alone cannot distinguish whether accurate judgments arise from continuously tracking physiological signals or from stable prior expectations about one’s heart rate **(Desmedt & Bergh, 2024)**. Distinguishing these alternatives requires explicitly modelling how perceptual beliefs respond to physiological fluctuations.

By analysing the behaviour of 518 participants performing the HRD task **(Legrand et al., 2022)**, we found that models assuming stable cardiac beliefs consistently outperformed models assuming trial-by-trial tracking of heart rate. This suggests that, during the timescale probed by the HRD task, perceptual decisions are generally guided more by stable internal beliefs than by ongoing cardiac afferents. This does not, however, imply that cardiac signals are absent from perception, but rather that their influence on behavioural reports is relatively limited under these experimental conditions.

This interpretation is further supported by a quasi-experimental analysis exploiting naturally occurring fluctuations in heart rate. Despite clear differences in physiological state across trials, inferred cardiac beliefs remained comparatively stable across individuals. These findings converge with growing evidence that behavioural performance in cardiac interoception tasks often reflects stable physiological expectations in addition to sensory evidence **(Desmedt et al., 2018; Ferentzi et al., 2025; Körmendi et al., 2023; Palmer et al., 2025)**.

At the same time, belief updating was not uniformly absent. Bayesian updating models provided a better account of behaviour for about 40% of participants. Rather than treating interoceptive perception as either static or dynamic, our results support a continuous view in which individuals differ in the weight assigned to afferent evidence. Consequently, conventional measures of interoceptive accuracy, including HRD bias **(Banellis et al., 2026a; Courtin et al., 2025; Tyrer et al., 2025)**, should not be interpreted as direct measures of physiological tracking without carefully considering the dynamic contribution of prior beliefs. This observation further motivates our definition of cardiac interoceptive sensitivity as a computational measure of belief updating.

### Bayesian belief updating and interoceptive sensitivity

The idea that interoceptive perception depends on integrating priors with incoming physiological signals has been approached multiple times **(Brand et al., 2024; Epstein & Stein, 1974; Khalsa et al., 2009; Larsson et al., 2021; Mandler & Kahn, 1960; Pennebaker et al., 1982; Ponzo et al., 2021)**. Yet existing behavioural measures have lacked a principled way to quantify the relative contribution of these two sources of information, which we have captured here as interoceptive sensitivity.

We developed two Bayesian models to estimate this quantity, which only differed in whether perceptual priors remained fixed or evolved across trials. Despite their different assumptions, both models converged on the same conclusion: exteroceptive decisions relied predominantly on incoming sensory evidence, whereas interoception was more influenced by prior beliefs. This asymmetry was considerably larger than suggested by conventional psychophysical measures such as threshold or slope alone (see Section 11.1 for an overview of the estimation biases expected under each modelling assumption), indicating that modelling belief dynamics reveals computational properties that are not directly captured by standard behavioural parameters.

An important advantage of this framework is that physiological volatility becomes an informative signal rather than a nuisance variable **(Stephan et al., 2016)**. Rather than treating spontaneous fluctuations as noise, our models exploit them to estimate how strongly perceptual beliefs respond. The same logic underlies the estimation of homeostatic belief precision from regulatory signals, where the variance rather than the mean of the action sequence carries the identifiable information **(Unal et al., 2021)**. This perspective may generalise beyond cardiac interoception to other domains of embodied psychophysics in which the relevant sensory signals cannot be experimentally controlled.

At the same time, interoceptive sensitivity should not be interpreted as a pure measure of perceptual ability. The extent to which beliefs are updated necessarily depends both on the quality of sensory representations and on the strength of the underlying physiological signal. As previously suggested **(Desmedt et al., 2023)**, individuals may appear less sensitive because afferent signals are weak rather than because perceptual inference is impaired. Future models could therefore benefit from integrating richer physiological measurements across the autonomic hierarchy together with computational models of brain–heart interactions **(Candia-Rivera et al., 2022; Candia-Rivera et al., 2024; Unal et al., 2021)**, allowing perceptual sensitivity to be interpreted in the context of the physiological information available to the observer.

### Model evaluation and reliability

As to which method yields the most meaningful information, our results have highlighted a complex pattern. No single model dominates across evaluations, but all revealed complementary strengths. Static beliefs provide the most parsimonious account of behaviour. Cardiac belief is the most reliable latent estimate across relevant models (only dynamic-belief model does not estimate it), whereas Bayesian updating models recover computational quantities that more clearly differentiate interoceptive from exteroceptive cognition.

This distinction illustrates a broader trade-off in computational modelling. Simpler models may provide more robust estimates of stable behavioural characteristics, whereas richer generative models can reveal latent computational processes that are inaccessible to simpler approaches but may be estimated less reliably **(Hess et al., 2025; Wilson & Collins, 2019)**. Importantly, the Bayesian models were identified as the best-explaining model for 40% of the population, and both estimated cardiac beliefs were comparable to what the static belief elicited, which appears to provide both predictive and explanatory advantages.

The cardiac hierarchical Gaussian filter represents the most expressive model in this comparison. It provided the clearest computational separation between interoceptive and exteroceptive processing, indicating that the two modalities rely on markedly different balances between prior expectations and sensory evidence **(Toussaint et al., 2024)**. This increased discriminability, however, came at the cost of reduced reliability of individual sensitivity estimates, which appeared more difficult to infer in this setup.

### Limitations and future perspectives

Overall, our findings challenge the use of bias and correspondence as primary measures of cardiac interoceptive accuracy. Interoceptive sensitivity relies on time-resolved updating that would be conflated by traditional psychophysical models such as static and dynamic beliefs. Our models can be applied retrospectively to existing datasets, facilitating cumulative progress without requiring new data collection. More generally, our results do not challenge the experimental design of the HRD task itself as they target modelling decisions that are made to interpret the data.

However, it has been pointed out that this paradigm was optimised for measuring how changes in cardiac activity are detected and interpreted **(Adamic et al., 2025; Körmendi & Ferentzi, 2023)**. While we have argued with the pseudo-experiment design above that the rather short trials used in the experiment can be leveraged to approach time-dependent changes in the physiological signal, even a posteriori, some components such as the psychometric procedure are still built from different perspectives. The task **(Legrand et al., 2022)** uses Psi, a Bayesian staircase **(Kontsevich & Tyler, 1999)** to adjust next trial intensity via entropy minimization. This minimisation process uses uncertainties around the estimate of the threshold and slope, but other parameters could be included as well, such as the interoceptive sensitivity *λ*, either in its time-dependent or independent version. The psychometric procedure could itself be adapted to track a metric whose validity is less debated, such as cardiac belief or interoceptive sensitivity. However, as no method could a priori indicate which model or strategy a given participant would rely on most, this choice will always be debatable.

The critical information that could disentangle this is the observation of decisions under out-of-bounds cardiac frequencies. But critically, because sensitivity depends on physiological variability, psychophysical methods that discard out-of-distribution trials **(Kingdom & Prins, 2016)**, such as lapse-rate modelling **(Banellis et al., 2026a; Banellis et al., 2026b; Courtin et al., 2025; Tyrer et al., 2025)**, must be applied cautiously. Here, it introduces a chance that the model will disregard trials carrying most information about interoceptive sensitivity. The direct consequence is that this approach will induce optimistic biases towards a precise slope (see also Section 11.1 for details on this point).

For that reason, our framework illustrates how naturally occurring physiological variability can replace experimental control when estimating latent perceptual processes, providing a computational foundation for embodied psychophysics. A notable limitation of our approach is the reliance on trial-averaged estimates of heart rate instead of beat-to-beat intervals. Because this smoothing over several systolic intervals (usually 4-6) likely limits temporal precision, access to full electrocardiographic recordings would enable modelling belief volatility and time-varying sensitivity at finer scales. While the cardiac HGF presented here avoids volatility estimates (see Section 11.2), we could presume that the irregular and nonlinear patterns of heart rate variability might require variation in learning rate and dynamic patterns of cardiac sensitivity. Although the field of cardiac interoception has undergone phases of conceptual uncertainty **(Desmedt & Bergh, 2024)**, accompanied by growing concern over the limited clinical relevance of existing metrics **(Adams et al., 2022; Banellis et al., 2026a; Banellis et al., 2026b; Shamsabad et al., 2025)**, we propose that computational modelling of embodied psychophysics offers a crucial path forward both for metrics and task development. Integrating such data with state-space **(Rosas et al., 2023; Rosas et al., 2024)** or hierarchical models of cardiac dynamics represents a promising direction for future research in embodied psychophysics **(Brener & Ring, 2016)**.

## Conclusion

We introduced a computational framework that distinguishes stable cardiac beliefs from their dynamic updating by physiological afferents. Across the largest HRD dataset analysed to date, cardiac beliefs in healthy participants were only weakly influenced by moment-to-moment physiological fluctuations, despite substantial variability in heart rate. More broadly, our work demonstrates how naturally occurring bodily variability can be incorporated into generative models to estimate latent perceptual dynamics when sensory signals cannot be experimentally controlled. We anticipate that this framework will facilitate the development of mechanistically grounded measures of interoception and provide a foundation for future work in embodied psychophysics.

## Author contributions

**N.L**.: Conceptualization, Data curation, Formal analysis, Investigation, Methodology, Software, Visualization, Writing – original draft, Writing – review & editing. **L.W**.: Formal analysis, Methodology, Writing – review & editing. **C.M**.: Formal analysis, Methodology, Writing – review & editing.

## Data availability

The trial-level behavioural data analysed here are those published in **Legrand et al. (2022)** and **Banellis et al. (2026a)**. Deidentified data and scripts for the second cohort are archived at https://doi.org/10.5281/zenodo.18006899 and at https://github.com/embodied-computation-group/multi_intero. Preprocessed data and scripts for the first cohort are available at https://github.com/embodied-computation-group/CardioceptionPaper.

## Code availability

All code used to fit the models, run the simulations, and produce the figures is available at https://github.com/LegrandNico/ComputationalCardioception.

## Appendix

### Heart rate variability biases estimates of cardiac beliefs

The heart rate discrimination task estimates the bias and the precision of cardiac beliefs **(Legrand et al., 2022)**. Those estimates are consistent (i.e., they converge on the true values as trials accumulate) under two assumptions: that the reference signal is invariant 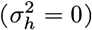, and that participants access it without bias. But these prerequisites are not met by human physiology and behaviors. Heart rate varies substantially within a session, is neither stationary nor independent across trials, and we cannot assume a priori that participants track those fluctuations rather than relying on a stable prior. When the reference moves and the fitted model assumes it does not, both the threshold and the slope of the psychometric function are biased.

We make that dependency explicit below for the two limiting strategies of the main text: the cardiac interoceptor, whose belief follows the heart on every trial (Model 1), and the cardiac believer, whose belief ignores it (Model 2). The result is symmetric: each strategy is recovered without bias in one coordinate system, and inflated by exactly 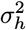 in the other. We describe both models explicitly below to further motivate the generative models used in this paper and discuss why possible correction methods should be applied carefully.

### Setup

The heart rate *h* recorded on trial *t* is a Gaussian random variable,

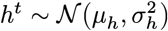

Similarly, cardiac beliefs are normally distributed over the same frequency continuum,

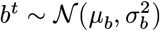

On an interoceptive trial, the staircase generates a comparison tone at an offset from the heart rate, *θ*_*i*_ = *h*_*t*_ + Δ_*t*_. Because the staircase tracks Δ rather than *θ*, the intensity Δ_*t*_ is independent of *h*_*t*_ by construction. Two coordinate systems are therefore available for the psychometric fit: the **relative** one, in Δ_*t*_, used by the dynamic-belief model, and the **absolute** one, in *θ*_*i*_, used by the static-belief model. Everything below follows from a single identity: marginalising a cumulative normal over a normally distributed mean returns a cumulative normal whose variance is the sum.

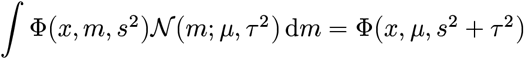

Unmodelled variability in the reference signal can therefore inflate an estimated variance by exactly the variance it contributes.

### Cardiac interoceptors

Cardiac interoceptors update their belief linearly under afferent signals, so the belief mean tracks the heart rate up to a bias *b*_*t*_ = *h*_*t*_ + *α*. The probability *p* of responding “Faster” on trial *t* is

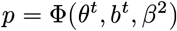

where the cardiac belief is conditioned on heart rate and bias

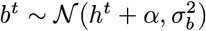

with 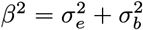 combines auditory and cardiac uncertainty.

Psychometric parameters using this model are

- **Unbiased in relative coordinates**. A psychometric fit in Δ_*t*_ recovers the threshold *α* and the variance B^2^ directly, regardless of heart rate variability.
- **Biased in absolute coordinates**. Reading the same behaviour as a belief over absolute frequencies requires marginalising over *h*_*t*_

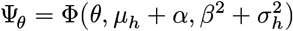

The inferred belief is centred on the mean heart rate rather than the trial-wise one, and its variance is inflated by 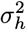: interoceptors read in absolute coordinates look less precise than they are.

### Cardiac believers

Cardiac believers compare the tone to a fixed internal reference. The trial-wise distance between tone and belief varies only because the heart rate under it moves. The probability of responding “Faster” is

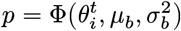

Psychometric parameters using this model are

- **Unbiased in absolute coordinates**. A psychometric fit in *θ*_*i*_ recovers *μ*_*b*_ and 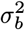 directly.
- **Biased in relative coordinates**. Using *θ*_*i*_ = *h*_*t*_ + Δ_*t*_ and marinalising over *h*_*t*_ yields

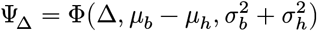

The threshold now measures the distance between the belief and the average heart rate. Under this model, the negative interoceptive threshold characteristic of the HRD task is not an underestimation of heart rate, and the variance is again inflated by 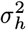.

### The interoceptive sensitivity continuum

For a participant who weights afferent evidence by *r* ∈ [0, 1], as in Models 3 and 4, the belief mean is 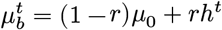, and the same marginalisation gives

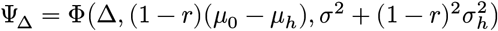

recovering the interoceptor at *r* = 1 and the believer at *r* = 0. Both the threshold and the slope of a conventional psychometric fit are therefore functions of the same unknown weight *r*, of the participant’s own heart-rate statistics, and of the belief parameters.

### Bias corrections

Since 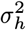 is observed, an obvious remedy would be to subtract it from the inferred belief precision.

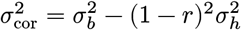

But *r* is the unknown of interest and, as we report here, exhibits large interindividual variability. Applying the correction should only be done after a robust estimation of this parameter. In this paper, we only considered modelling approaches that would allow robust inference of *r* and have not addressed correction methods for belief precisions or their performance.

However, we can see that other modelling choices could worsen biases in the precision estimates. Because the 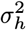 term inflates the tails of the psychometric function, procedures that attribute tail errors to attentional lapses **(Banellis et al., 2026a; Banellis et al., 2026b; Courtin et al., 2025)** remove exactly the variance that carries information about belief updating. Such procedures steepen the fitted slope and return an optimistically precise estimate of cardiac belief precision. For that reason, this should be applied cautiously to interoceptive data.

### Derivation of the sensitivity metric in the cardiac HGF

The perceptual sensitivity *λ* is given by:

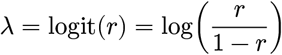

Where

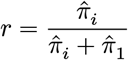

With *π*_*i*_ the precision of the input node and *π*_1_ the precision of the value parent. And we can derive the full sensitivity as:

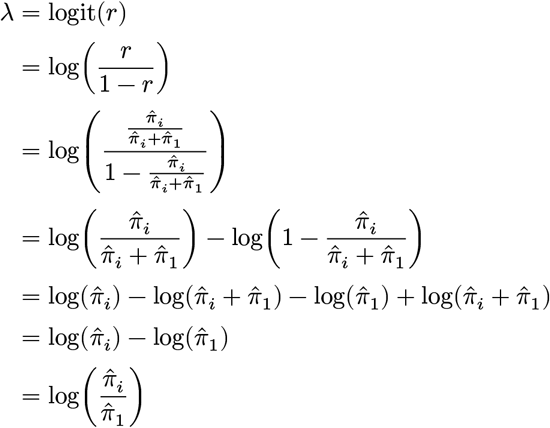

### Graphical models

We report below the graphical models used and reported by PyMC **(Abril-Pla et al., 2023)** to infer parameters for one participant. Readers who are interested in accessing the parametrization of the distribution are directed to the source code for the models, which can be accessed at https://github.com/LegrandNico/ComputationalCardioception/tree/main/code/models.

### Dynamic beliefs

**Figure 6:**
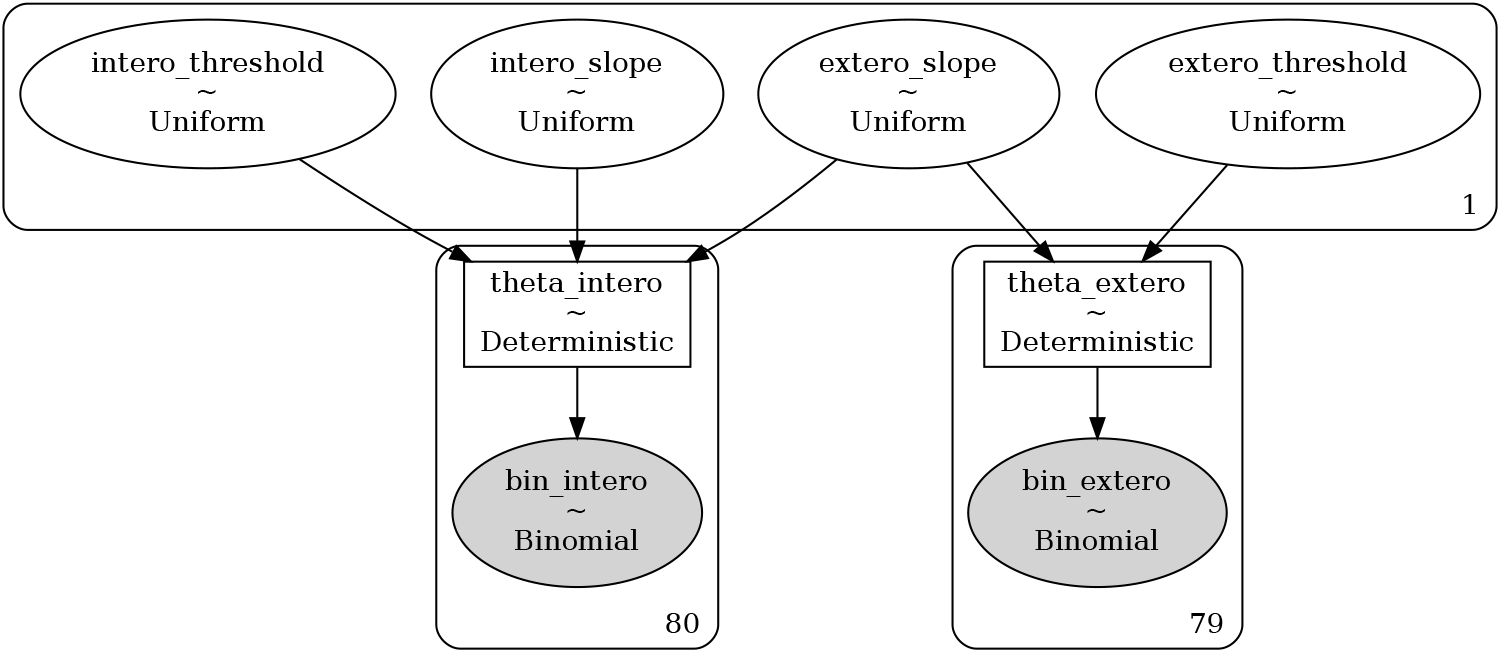
Graphical model used to fit the model of dynamic beliefs.

### Static beliefs

**Figure 7:**
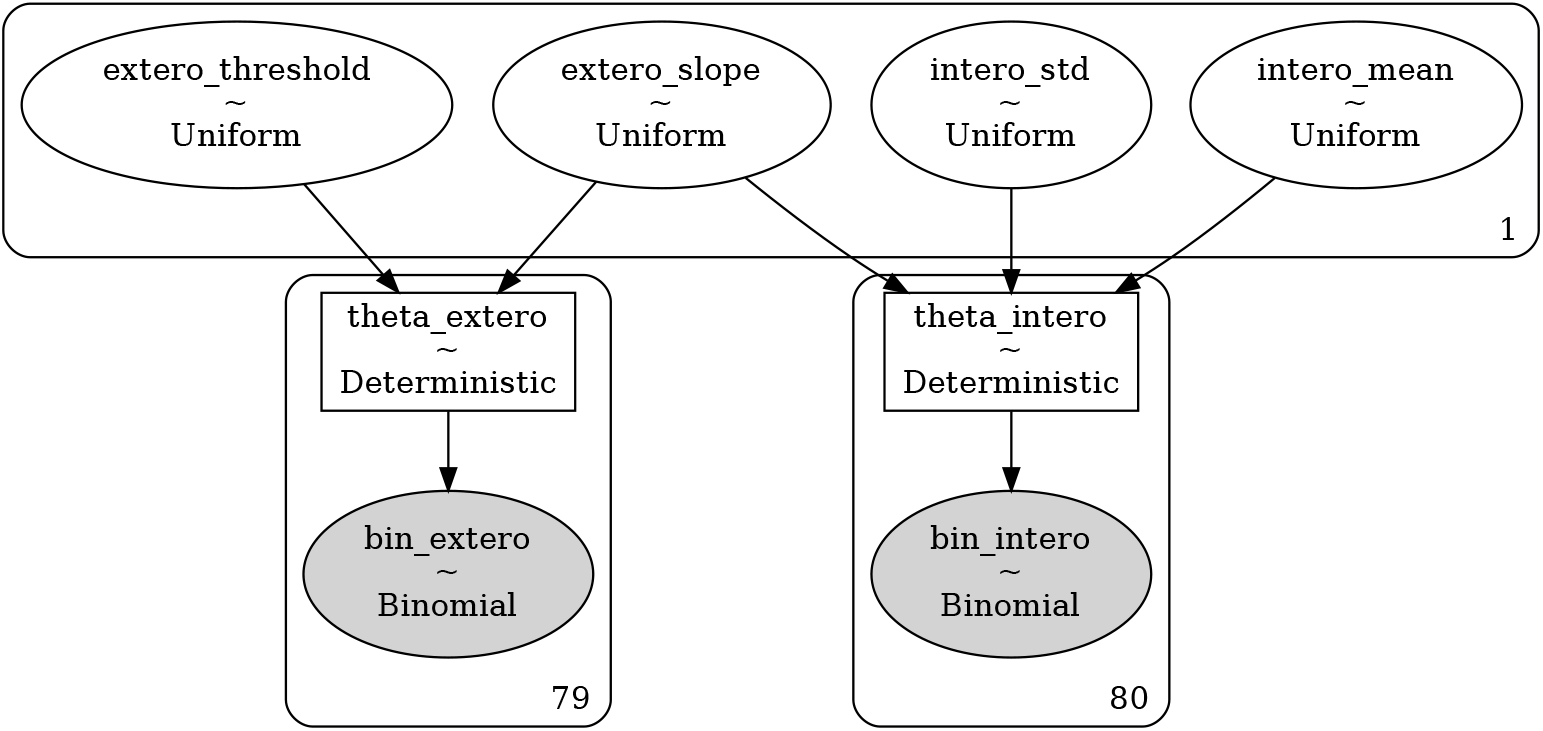
Graphical model used to fit the model of static beliefs.

### Weighted Bayesian update

**Figure 8:**
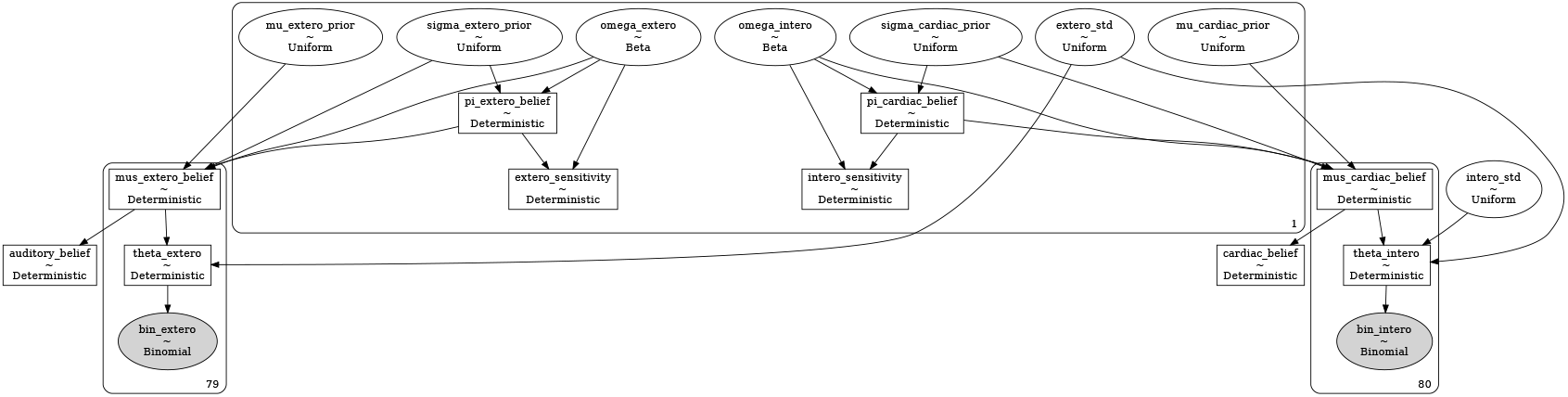
Graphical model used to fit the model of weighted Bayesian updates.

### Weighted Bayesian update with dynamic priors

**Figure 9:**
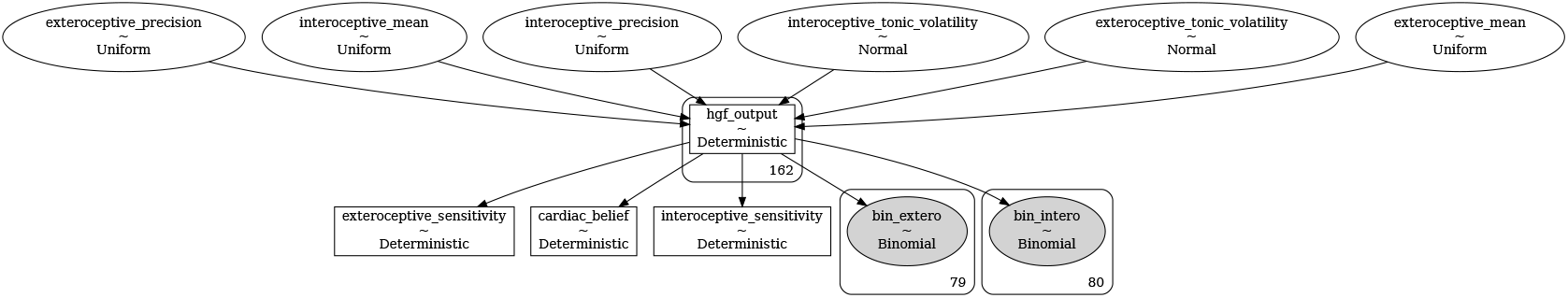
Graphical model used to fit the model of weighted Bayesian updates with dynamic priors (cardiac HGF).

### Model fit on a single participant

**Figure 10:**
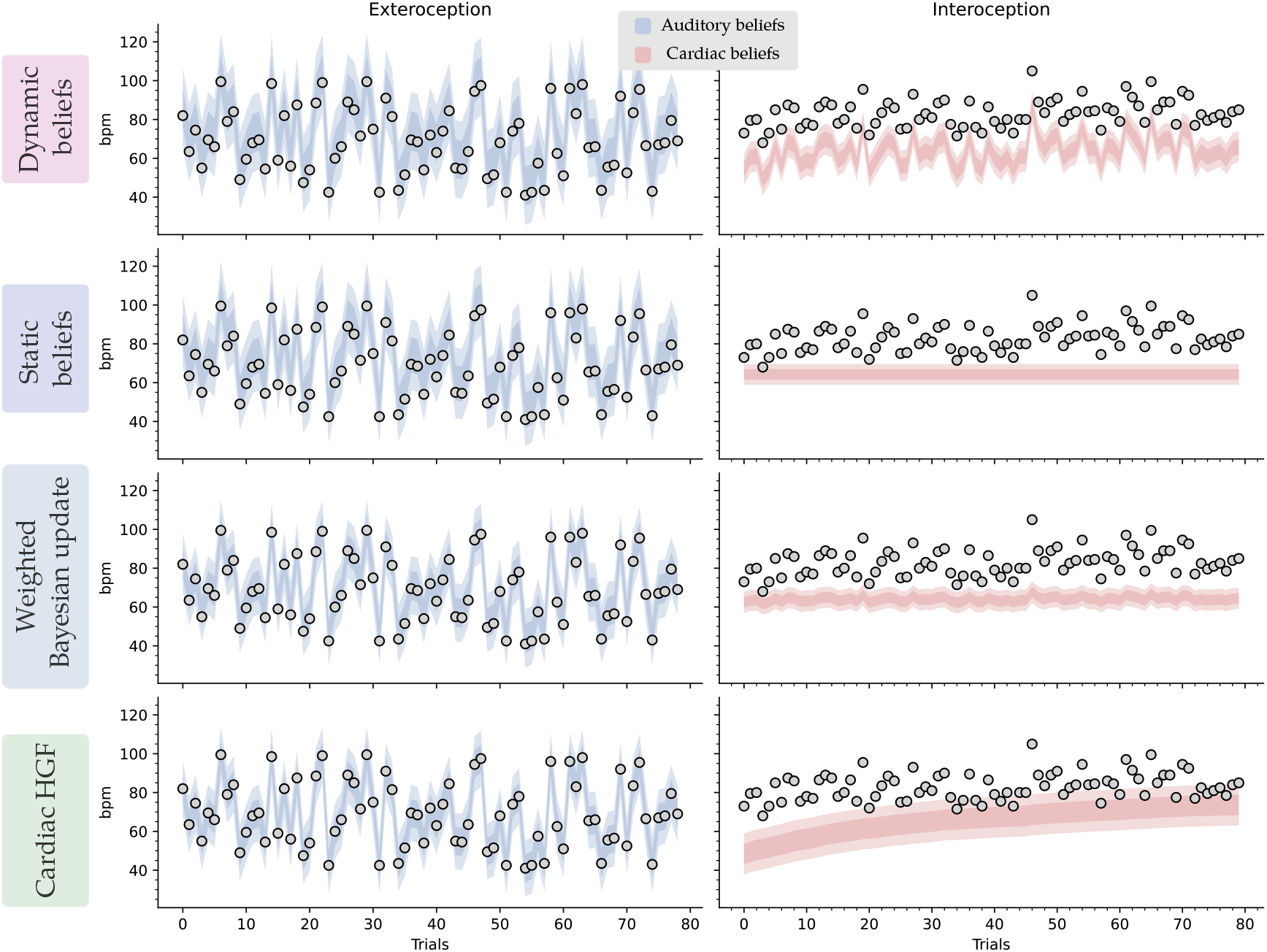
Example of model fit on a representative participant for exteroception and interoception trials. The shaded areas represent 1 and 2 standard deviations from the mean. The beliefs represent the inferred state of the first stimulus, either the first tone (exteroceptive trials) or the heart rate (interoceptive trials). Exteroceptive beliefs show larger volatility, as they are almost only influenced by the new stimulus and do not carry prior expectations. Cardiac beliefs, on the other hand, are more resistant to change, as attested by the slow evolution in the two update models (i.e., the Cardiac HGF and the weighted Bayesian). Importantly here, the model is fit to participants’ behaviors, which are influenced by beliefs, and as such it is expected in the interoceptive modality that the curve poorly fits the instantaneous heart rate.

